# Seeing is Feeling: How Aphantasia Alters Emotional Engagement with Stories

**DOI:** 10.1101/2025.01.09.632075

**Authors:** Noha Abdelrahman, David Melcher, Pablo Ripollés

## Abstract

Visual imagery is thought to act as an ‘emotional amplifier’, potentially contributing to narrative engagement. To examine this, we conducted two experiments in which participants were presented with emotionally charged audio and video story excerpts. Experiment 1 included 84 online participants from the general population, while Experiment 2 involved 25 individuals with aphantasia (the inability to generate mental images) and 25 controls. In both experiments we assessed narrative engagement behaviorally using the Narrative Engagement Questionnaire (NEQ), while for Experiment 2 we also measured physiological responses. We found a main effect of modality, with video stimuli scoring higher across all NEQ subscales in both experiments. Notably, in experiment 2, a significant group effect on emotional—but not cognitive—engagement emerged, with aphantasics reporting less emotional engagement than controls. Moreover, controls experienced higher heart rate during audio narratives, while aphantasics had a similar heart rate across both modalities. Our results suggest that the enhanced physiological response seen in non-aphantasics during audio narratives is driven by the mental effort required to generate imagery. Furthermore, this capacity for visual imagery appears to enhance emotional engagement with stories. This highlights mental imagery’s role in both subjective and physiological responses, emphasizing distinct cognitive processes during narrative engagement.

## 1. Introduction

Mental imagery (i.e., the ability to create mental images) occurs frequently during everyday life (Blackwell et al., 2020) and plays a key role in cognition, including in aspects of decision-making, planning actions, and emotion regulation (Pearson et al., 2015; Isaac and Marks, 1994), among others. Mental imagery is thought to act as an ‘emotional amplifier’ and this has significant real-life implications. For instance, mental imagery can enhance the treatment of PTSD by reducing the vividness of intrusive imagery (Holmes et al., 2008; Chuang, 2021; Morina et al., 2013), and it can aid in building healthy habits by increasing the vividness of future events and rewards, an intervention that can be used, for example, in smoking cessation programs (Stein et al., 2016; Bulley et al., 2019).

Mental imagery also modulates the way we engage and learn from narratives and stories (e.g., books: Boerma et al., 2016). Stories are powerful—they do not just captivate us; they can change us. Studies reveal that stories shape our beliefs and attitudes (Green et al., 2003; Marsh et al., 2003; Rapp, 2016; Gerrig, 1993; Quinlan & Mar, 2020), even our views on others (Mar, 2018). Koopman (2015, 2016) showed how fiction makes us reflect deeply, especially on challenging emotions. Gardner & Knowles (2008) found that fictional characters can impact us like real people through emotional bonds. Moreover, ‘bibliotherapy’—using stories for healing (Tribe et al., 2021; Troscianko, 2018; Hamm & Leonhardt, 2023)—highlights fiction’s therapeutic power. In this context, mental imagery helps to make stories potent tools for shaping and influencing us (Quinlan & Mar, 2020).

The role of mental imagery in narrative emotional engagement has been a prominent focus of psychological research, particularly within studies focusing on assessing a uniquely human skill: reading. This literature has explored mental imagery both as an individual trait (Denis, 1982; Long et al., 1989; Weibel et al., 2011; Isberner et al., 2019; Blazhenkova et al., 2023; Troscianko, 2013; Slámová, 2020; Wicken et al, 2021), and as as a state (imagery evoked during reading; Grossberg & Wilson, 1968; Nell, 1988; Sadoski, 1988; Sadoski, 1983; Irwin & Witte, 1980; Brück et al., 2016; Öncel et al., 2020; Johnson et al., 2013; Webster & Saucier, 2011; Webster & Saucier, 2011; Holmes & Mathews, 2005; Green et al., 2003). While several studies have suggested that visual mental imagery is necessary for emotional engagement with storytelling, others have found no such relationship (Nell, 1988; Green et al., 2008). To provide a more definitive answer to this question, a few studies have focused on exploring the association between mental imagery and narrative engagement within the two extremes of the vividness of the imagery spectrum: aphantasia (the inability to generate and experience mental imagery) and hyperphantasia (the ability to generate extremely vivid visual mental images; Zeman et al., 2020; Milton et al., 2021). However, these studies have also provided inconsistent results, with research both confirming the association (Speed et al., 2024; Wicken et al., 2021), and showing opposite patterns (Troscianko, 2013; Slámová, 2020).

Importantly, the degree to which imagery contributes to narrative engagement tends to vary depending on the format in which the story is delivered. Several authors have hypothesized that reading novels tends to engage one’s visual imagination to a greater extent than watching movies (Weibel et al., 2011; Isberner et al., 2019; Long et al., 1989; Jajdelska et al., 2019). In this vein, a recent study compared emotional engagement induced during video (e.g., movies, tv series, etc.) versus audio narratives (Richardson et al., 2020). Although participants self-reported heightened engagement during video presentations compared to auditory scenarios, the physiological data exhibited the opposite pattern, with auditory narratives triggering increased physiological responses (electrodermal activity, heart rate, etc.) as compared to video stimuli. Richardson and colleagues (2020) suggested that mental imagery generated in the listener’s mind may not attain the same vividness and detail during spoken narratives compared to their on-screen counterparts and are thus often explicitly rated as less engaging. However, the construction of the mental images from auditory narratives without visual stimulation necessitates more profound cognitive and emotional processing, rendering them more physiologically engaging. The challenge lies in discerning whether these physiological changes stem from cognitive effort, related to imagery construction, or emotional arousal, as both factors similarly influence the autonomic nervous system (Potter, 2011; Sequeira et al., 2009; Sukalla et al., 2015; Andreassi, 2006). This ambiguity suggests that observing heart rate or electrodermal activity alone may not clearly differentiate between cognitive and emotional influences. Moreover, Richardson et al. (2020) did not account for individual differences in mental imagery, which is the proposed key contributor to the mental effort involved in this context. This limitation complicates the interpretation of whether the observed physiological responses are driven by emotional arousal or the cognitive effort required to generate mental images.

Here, we aim to test the hypothesis that emotional engagement with storytelling is amplified by mental imagery capacity, and that increased physiological responses while listening to audiobooks as compared to watching videos stem from the additional mental effort required to generate mental images. To do so, we focused on a special population which reports a lack of visual mental imagery ability: aphantasics. Our objective was to provide a better understanding of mental imagery using aphantasics as a naturally occurring ‘knock-out’ model of imageless cognition (Keogh and Pearson, 2018). In a first online experiment (N=84), we attempted to replicate Richardson and colleague’s (2020) behavioral results in which participants from the general population reported video stories to be more emotionally engaging than audio ones. In a second experiment, we extended Richardson and colleagues’ (2020) methodology to include aphantasic participants (N=25) and a group of matched controls (N=25), while collecting behavioral and physiological measures (electrodermal activity and heart rate). We specifically predicted that aphantasic individuals, who lack the ability to create visual images, would experience reduced emotional engagement when listening to audio narratives compared to control participants with intact imagery. We further hypothesized that aphantasic individuals would not show an increased physiological response during audiobook listening, likely due to the absence of the additional cognitive load needed to create and sustain mental images that is present in control participants.

## 2. EXPERIMENT 1

### 2.1. Methods

#### 2.1.1. Participants

90 participants were recruited online on Prolific Academic (https://prolific.co/) and from NYU’s student population. Six participants were excluded due to technical failures, participant dropouts, or duplicate participation in the experiment, leaving a total sample of 84 native English speaker participants (age: 39 ± 15 (SD), 55% females). All participants provided online written informed consent and were paid for their participation. The experiment was approved by NYU’s IRB.

#### 2.1.2. Stimuli and materials

##### 2.1.2.1. Stories

The stimulus set consisted of six scenes from different stories, both in the form of audiobooks and videos. These stimuli were derived from the set created by Richardson and colleagues (2020). The authors curated a collection of fictional works that encompassed distinct genres. We used six stories from three different genres. To represent classic literature, *Pride and Prejudice* and *Great Expectations* were included. From the realm of crime fiction, *Hound of the Baskervilles* and *The Silence of the Lambs* were chosen. Lastly, from the Science Fiction/Fantasy genre, we included *Alien: River of Pain* and *A Song of Ice and Fire* (Game of Thrones). The audiobooks are faithful readings of the original texts, rather than adapted audio plays with additional sound effects and actors. The videos, on the other hand, were extracted from film or TV adaptations of the books. Each story was selected based on its recognition as a prominent example within its respective genre, as well as the availability of both audio and video adaptations. The scenes selected from each story were those that possessed emotional intensity and were suitable for presentation in both audio and video formats. Those scenes aimed to evoke strong emotional responses and maintain comparability between the two formats. While the events covered were the same, due to their inherent nature, the audiobooks tended to have longer durations (M = 392 secs, SD = 227) compared to the video versions (M = 298 secs, SD = 133). This difference, however, was not statistically significant (Wilcoxon Z = 1.153, p = 0.313). A detailed description of the six scenes can be found in the Supplemental Information.

#### 2.1.3. Design and Procedure

The experiment was coded using PsychoPy version 2021.2.4 (Peirce et al., 2019) and was hosted online on the Pavlovia server (https://pavlovia.org). Firstly, participants provided informed consent and completed a survey of demographic data (age, gender) through Qualtrics (https://www.qualtrics.com/). They were then redirected to Pavlovia to take the experiment.

Participants were presented with six scenes in total, presented in two blocks: three scenes were in the form of an audiobook, and the other three were in the form of videos. The order of these blocks, as well as which stories were presented in each format, were randomized across participants. Specifically, participants were randomly assigned to begin with either an audiobook or video block. Each block included three randomly chosen scenes. Once a scene was presented in one format (audiobook or video), it was not repeated in the opposite format. For instance, if a participant started with audiobooks and listened to “Alien”, “Pride and Prejudice”, and “Hound of Baskerville”, these stories would not appear again in the subsequent video block. Instead, they would see “Silence of the Lambs”, “Game of Thrones”, and “Great Expectations”. Each trial (for both videos and audiobooks) began with participants reading a concise summary of the story’s plot and characters thus far, providing context for the upcoming excerpt (Richardson et al., 2020). Subsequently, they either viewed the video onscreen or listened to the story while observing a white screen. Following the scene, participants used a slider (ranging from 1 to 5) to rate their familiarity with the story. Then they completed the Narrative Engagement Questionnaire (NEQ) by Busselle and Bilandzic (2009), with the questions presented in a randomized order. The questionnaire captures different aspects of engagement and includes 4 subscales. These subscales capture aspects of engagement related to cognitive processing such as narrative understanding (e.g., “At points, I had a hard time making sense of what was going on in the story”) and attentional focus (e.g., “I found my mind wandering while the story was playing”), and others related to emotional engagement such as narrative presence (e.g., “During the story, my body was in the room, but my mind was inside the world created by the story”) and character engagement (e.g., “I felt sorry for some of the characters in the program”). Participants evaluated the 12 statements (3 per subscale) of the questionnaire utilizing a 7-point Likert scale ranging from “strongly disagree” to “strongly agree”. As in Richardson and colleagues (2020), each subscale was scored by summing their 3 corresponding questions, giving a value between 3 and 21. The experiment lasted about an hour, after which participants were compensated for their participation.

#### 2.1.4. Statistical Analyses

To analyze narrative engagement as measured with the NEQ, we conducted linear mixed modeling using R (version 4.0.2) and RStudio (version 1.3.959; Bates et al., 2015), employing the *lme4* package. Four models were calculated, using each of the four NEQ subscale scores as dependent variables. All the models included modality (audio vs. video) and familiarity as fixed effects, and an interaction between these two factors. We also included random intercepts for each participant and each story.

To evaluate the impact of various predictors, we utilized the *car* and *emmeans* packages in R, conducting Type III Wald chi-square tests.

### 2.2. Results

We observed a significant main effect of modality on each of the four dimensions of narrative engagement, replicating Richardson and colleagues’ (2020) behavioral results (Figure 1; Table 1). Participants reported greater narrative presence, engagement with characters, attentional focus, and understanding when experiencing video narratives compared to audio narratives (p < 0.001 for each scale; see Figure 1A).

**Figure 1.**
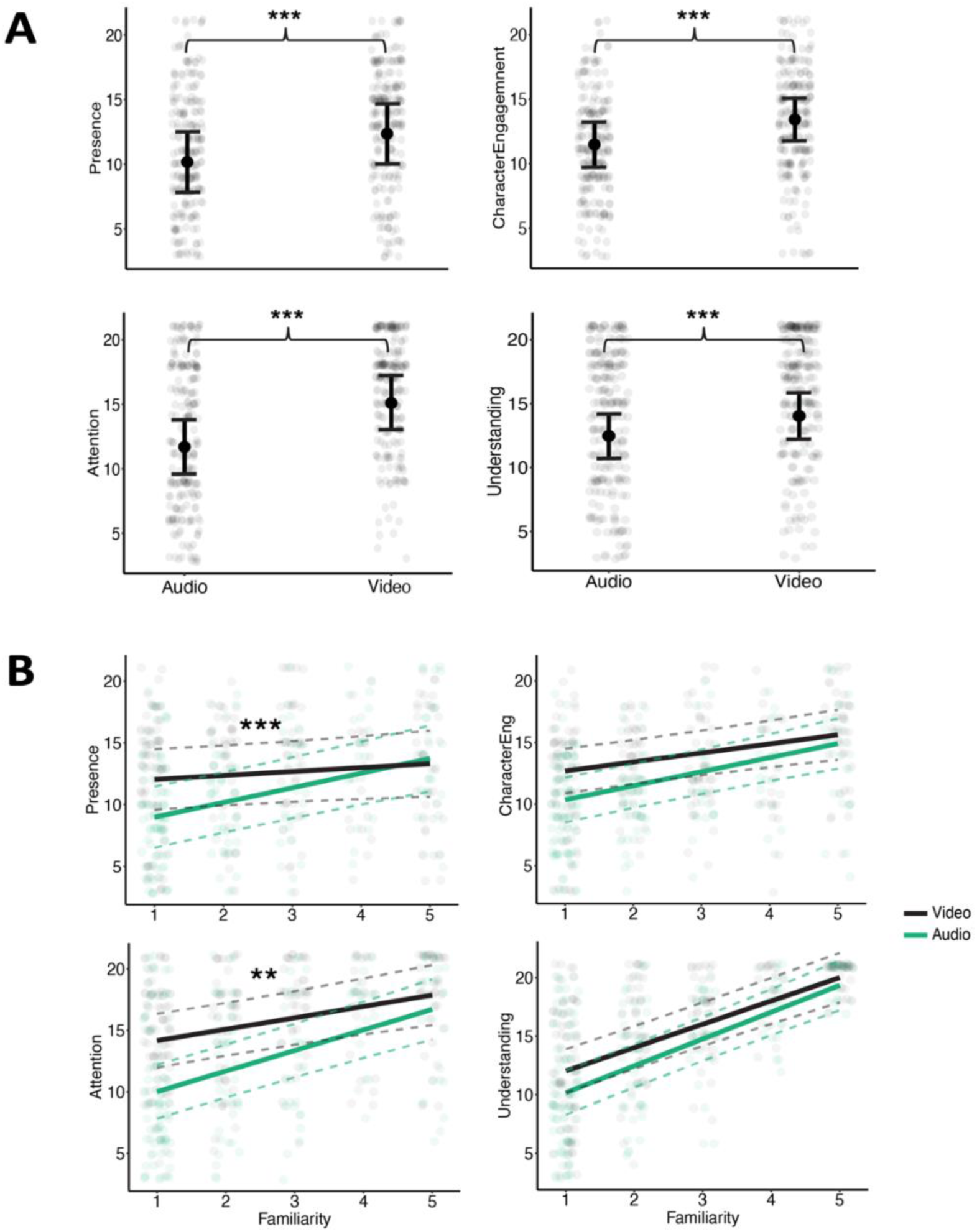
Experiment 1. (A) Model-predicted means and 95% confidence intervals of the narrative engagement questionnaire subscales split between story modalities. (B) Correlations of familiarity with the narrative engagement questionnaire subscales split between story modalities. Note that we have marked only the effects driving interactions (presence and attention); we have not highlighted the familiarity main effects. ***, p<0.001. **, p<0.01

**Table 1.**
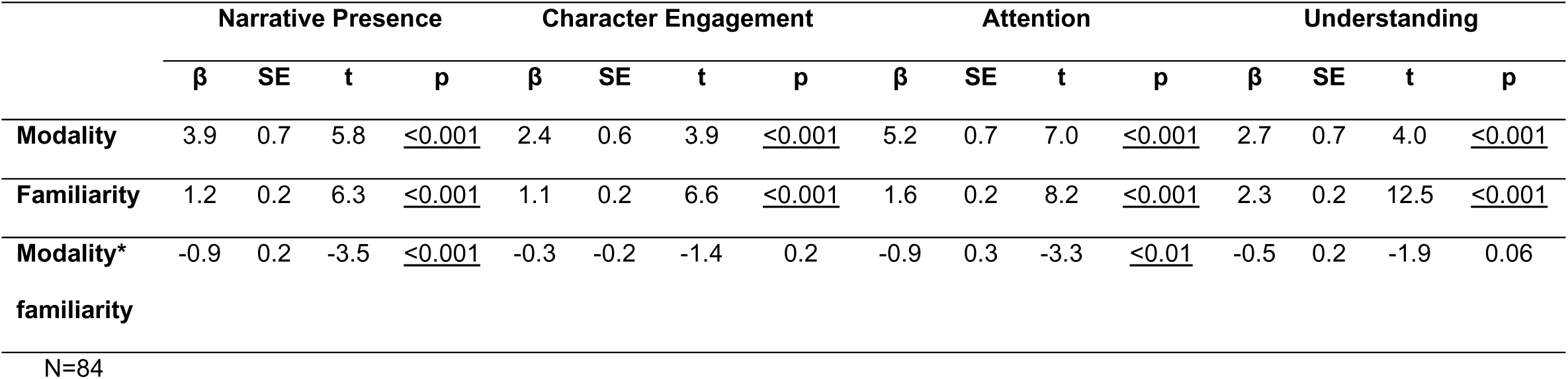
Linear mixed models predicting narrative presence, character engagement, attentional focus and understanding.

There was also a significant main effect of familiarity on each of the four subscales (i.e., the more familiar the stimulus, the greater the engagement; p < 0.001; see Figure 1B). Additionally, the interaction between modality and familiarity was significant for narrative presence and attentional focus (i.e., the slope describing the relationship between engagement and familiarity is steeper for audio than for video; p < 0.001 and p < 0.01, respectively), but not for character engagement and understanding (p = 0.2 and p = 0.06, respectively). The results of the Linear Mixed Models (LMMs) exploring each subscale are detailed in Table 1. In summary, our findings indicate that video narratives consistently elicited higher engagement across all measured dimensions compared to audio. Familiarity further enhanced engagement, particularly within the audio format, where it significantly boosted narrative presence and attentional focus.

## 3. EXPERIMENT 2

To further assess the role of visual imagery in narrative engagement, we conducted a second, in-person experiment. This phase included physiological response measurements alongside the behavioral measures, and expanded our methodology to incorporate both aphantasic participants and control participants. Aphantasia served as a naturally occurring ‘knock-out’ model for imageless cognition, providing insights into the cognitive processes underpinning narrative engagement.

### 3.1. Methods

#### 3.1.1. Participants

In this in-person study, a total of 25 aphantasic participants (age: 35 ± 15 SD, 11 females) were recruited via Reddit (https://www.reddit.com), the Aphantasia Network (https://aphantasia.com/), and word of mouth. All aphantasic participants were native English speakers, except for one individual who, despite not being native, was highly fluent due to having lived in the United States for 20 years. In addition, 25 control participants (age: 33 ± 13 SD, 13 females) were recruited through the NYU paid research participation system and word of mouth. Five trials from four different participants were excluded due to technical issues or interruptions, such as participants speaking during the trial. All participants provided informed consent. The experiment was approved by NYU’s IRB. All participants in the aphantasia group scored below 32 on the VVIQ (min=16, max=31; mean 18 ± 3 SD). On the other hand, all participants in the control group scored above 32 (min=41, max=77; mean 60 ± 10 SD), consistent with previous studies (Zeman et al., 2015; Tabi et al., 2022). As expected, VVIQ scores were significantly higher in the control group than in the aphantasic group (t(48)=-19.7, p>0.001). To compare age and gender distribution between the two groups we used independent samples t-tests and Bayes factors (BF) computed in JASP using default priors (Morey et al., 2024; Rouder and Morey, 2012). We report BF_01_, which reflects how likely data is to favor the null hypothesis (i.e. the probability of the data given H0 relative to H1). There were no significant differences in terms of age and gender between the control and the aphantasic group (Age: t(48)=0.5,p=0.6, BF_01_=3.2; Gender: *X^2^*(1, N=50)=0.3,p=0.6, BF_01_=2.5)

#### 3.1.2. Stimuli and materials

The same stimuli set and materials were used as in Experiment 1 (Section 2.1.2.), with the addition of the Vividness of Mental Imagery Questionnaire (VVIQ: Marks, 1973) and the Aesthetic Responsiveness Assessment (Schlotz, 2021).

The VVIQ is a widely used self-report measure designed by Marks (1973) to assess an individual’s subjective experience of visual mental imagery. The questionnaire requires the participants to visualize four different scenes of different people, objects, and settings in their mind (e.g., “Think of the front of a shop to which you often go”). They are then asked to rate how clearly or vividly they were able to form those mental images. Scores on the VVIQ range from 16 to 80. The cut-off score for aphantasics is usually 32 (Dance et al., 2021; Dance et al., 2021; Wicken et al., 2021), which we also adopted in this study. This means that the participant gives at most a rating of 2 (vague and dim) out of 5 (perfectly clear and as vivid as normal vision) on average for the 16 questions. Some authors have used a stricter score of <25 (Bainbridge et al., 2021) and <23 (Zeman et al., 2020). A participant is considered a hyperphantasic if the score is > 75 (Zeman et al., 2020).

The Aesthetic Responsiveness Assessment (AReA) focuses on assessing typical responses to and engagement with various aesthetic stimuli, with a particular emphasis on visual aesthetic experiences. It uses 14 items, with statements like “I am deeply moved when I see art,” to assess participants’ levels of aesthetic appreciation, intense aesthetic experience, and creative behavior.

#### 3.1.3. Design and Procedure

After completing the same task as in Experiment 1 (Section 2.1.3.; i.e., experiencing the whole six stories and providing the behavioral ratings), participants completed the VVIQ. For the VVIQ, they indicated the level of vividness of their mental images using a 5-point Likert scale that ranged from ‘no image at all’ to ‘exactly clear and as vivid as regular vision’. They then rated each item of the AReA on a five-point scale from 1 ‘Never’ to 5 ‘Very Often’. Additionally, participants used two 5-point sliders to self-report their reading habits: the first asked them to rate ‘How good a reader are you?’ with options ranging from ‘Poor’ to ‘Excellent,’ while the second asked them to indicate how many novels they read per year, with options ranging from ‘Less than 4’ to ‘More than 16’. Participants were situated within a semi darkened room in front of a Dell windows desktop computer monitor at a distance of approximately 60 centimeters, and wore headphones throughout. Physiological data was measured using a Biopac System consisting of the acquisition module MP160 and collected at a sampling rate of 2000 Hz using the AcqKnowledge 5.0.1 software (Biopac Systems Inc., Goleta, CA, United States). Electrodermal activity (EDA) was measured using finger electrodes (EL507A). The electrodes were affixed using isotonic gel (GEL101A) and positioned on the middle and index fingers of the left hand. Participants were asked to provide responses with their right hand, which was electrode-free. These electrodes were connected to an EDA amplifier (EDA100C). Electrocardiogram (ECG, i.e., heart rate) measurements were taken using the ECG100C amplifier, with electrodes positioned in a lead-III configuration on the right collarbone and the lower ribs. Participants were instructed to keep still as much as possible throughout physiological data collection, especially when a story was being played.

#### 3.1.4. Physiological data preprocessing

EDA responses for each narrative were computed through the MATLAB EDA toolbox for psychophysiology data management, cleaning and analysis (Joffily, 2018; Ibáñez et al., 2016; Jaber-López et al., 2014; Sklivanioti Greenfield et al., 2023; Abrams et al., 2024). Initially, the EDA signal was isolated for each trial and preprocessed using a Butterworth low-pass filter with a cutoff frequency set at 1Hz, followed by downsampling to 100Hz. From this signal, we calculated phasic and tonic EDA responses. For phasic EDA, skin conductance responses exceeding 0.02 μs within a trial were detected automatically using the ‘eda_edr’ function. For all participants and all trials, the automatic detection was reviewed and manually corrected when needed. During this manual correction, we removed the first major skin conductance response in each trial, as it is usually related to a startle reflex driven by the sudden start of the sound (Bari et al., 2018). The base-to-peak difference of all identified phasic EDA responses per trial were added and then normalized by the duration of the narrative in minutes to account for variations in story length, then finally transformed into z-scores. EDA tonic response was calculated as the mean of the signal for each trial (Braithwaite et al., 2013) and then transformed into z-scores.

The raw ECG (i.e., heart rate) signals were detrended, and the heartbeats were identified as the R wave maxima of the QRS complexes. Custom code was used to remove noise, followed by a visual inspection of the ECG signals to ensure accurate identification of the QRS complexes (Palumbo et al., 2024). One aphantasic participant was excluded from the ECG analyses due to an insufficient signal-to-noise ratio. Another two control participants were excluded due to heart related medical conditions. Heart rate was calculated by dividing the sampling rate (samples per second) by the average number of samples per beat and then multiplying by 60 seconds per minute (Gorr, 2023), then finally transformed into z-scores.

#### 3.1.5. Statistical Analyses

To analyze narrative engagement as measured with the NEQ, we conducted linear mixed modeling using R (version 4.0.2) and RStudio (version 1.3.959; Bates et al., 2015), employing the lme4 package. Four models were calculated each using the NEQ subscale scores as dependent variables. Physiological data was analyzed using the same methodology. In particular, three models were computed, using, heart rate, tonic EDA and phasic EDA as dependent variables. All models (behavioral and physiological) included fixed effects for modality (audio vs. video), group (aphantasics vs. controls), and familiarity, as well as an interaction term between modality and group. We incorporated random intercepts for each participant and each story in all models. To evaluate the impact of various predictors, we utilized the *car* and *emmeans* package in R, conducting Type III Wald chi-square tests. Post-hoc tests were used to further assess significant effects and interactions. The resulting p values from these analyses were corrected for multiple comparisons using the Tukey method.

To assess differences in reading habits and aesthetic responsiveness between groups we used independent samples t-tests and Bayes Factors.

### 3.2. Results

#### 3.2.1. Behavioral Results

Replicating the results of Experiment 1 (Section 2.2.) and Richardson et al. (2020), we found that video narratives elicited greater narrative engagement than audio stories across all four subscales (p < 0.001 for each) regardless of the group. Familiarity also had a significant effect on each of the four subscales (p < 0.001). Detailed findings of the LMMs exploring each dimension can be found in Table 2 and Figure 2.

**Figure 2.**
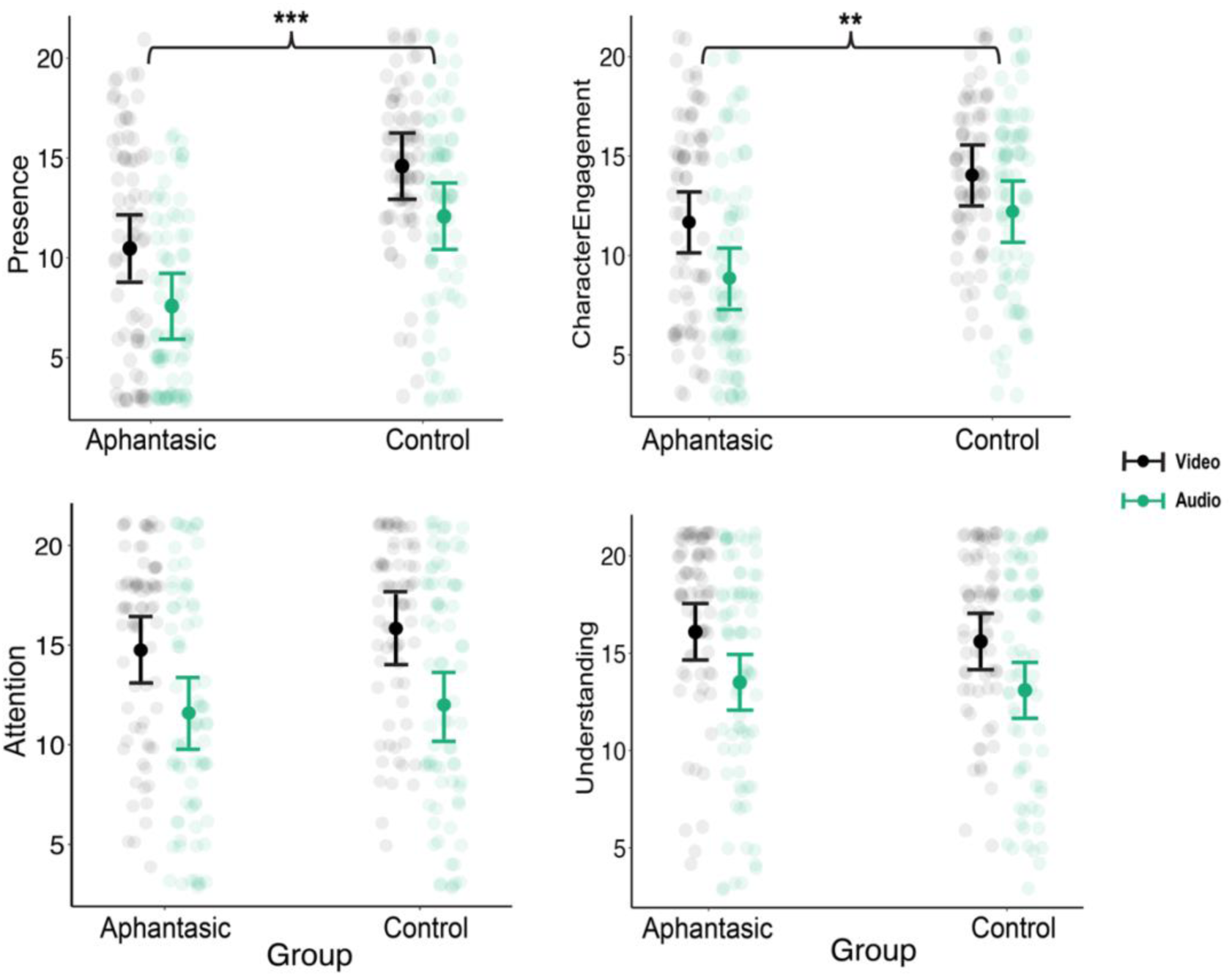
Experiment 2. Model-predicted means and 95% confidence intervals of the different dimensions of the self-reported engagement split between story modalities (audio vs video) and groups (controls vs aphantasics). Note that we have marked only the significant results driven by group effects (presence and character engagement); we have not highlighted the modality effects. ***, p<0.001. **, p<0.01. *, p<0.05

**Table 2.**
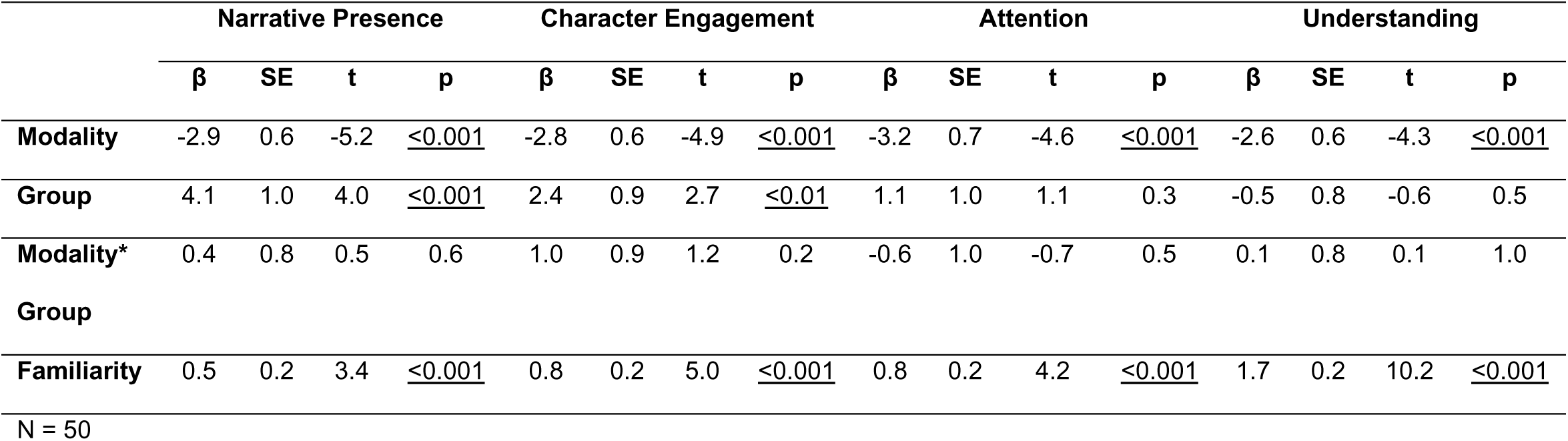
Linear mixed models predicting narrative presence, character engagement, attentional focus and understanding.

##### 3.2.1.1. Emotional subscales: Presence & Character Engagement

The analysis revealed a significant main effect of group on both presence and character engagement (p < 0.001 for both). Aphantasics experienced a lower sense of presence compared to controls while listening to audiobooks (β = -4.5, SE = 1.0, p < 0.001) and while watching videos (β = -4.1, SE = 1.0, p < 0.01). A similar pattern was observed for character engagement, where aphantasics rated their engagement significantly lower than controls for both audio (β = -3.4, SE = 0.9, p < 0.01) and video (β = -2.4, SE = 0.9, p < 0.05). Furthermore, we found no significant interactions between group and modality for neither character engagement (p = 0.2) nor presence (p = 0.6).

##### 3.2.1.2. Cognitive subscales: Attentional Focus & Narrative Understanding

We did not find a significant effect of group on neither attentional focus (p = 0.3) nor narrative understanding (p = 0.5). There was also no significant interaction between group and modality for neither attentional focus (p = 0.5) nor narrative understanding (p = 1.0).

##### 3.2.1.3. Reading habits

No significant differences were found regarding the perceived reading ability between aphantasics (mean = 3.8, SD = 1.1) and controls (mean = 4.0, SD = 1.0; t(48) = -0.5, p = 0.6, BF_01_ = 3.1). In terms of the number of novels read annually, aphantasics reported reading more novels (mean = 2.8, sd = 1.5) than controls (mean = 2.2, sd = 1.3), but the difference was not significant (t(48) = 1.6, p = 0.1, BF_01_ = 1.3).

##### 3.2.1.4. Aesthetic responsiveness

No significant differences were found regarding aesthetic responsiveness between aphantasics (mean = 2.8, SD = 0.7) and controls (mean = 3.1, SD = 0.5; t(48) = -1.4, p = 0.2, BF_01_ = 1.6).

#### 3.2.2. Physiological Results

##### 3.2.2.1. EDA

No significant effects were observed for phasic EDA in terms of familiarity (p = 0.1), group (p = 0.5), modality (p = 0.1), or the interaction between group and modality (p = 0.3; see supplemental Figure 1A). Similarly, no significant effects were found for tonic EDA concerning familiarity (p = 0.6), group (p = 0.4), modality (p = 0.1), or the interaction between group and modality (p = 0.2; see Table 3 and supplemental Figure 1B).

**Table 3.**
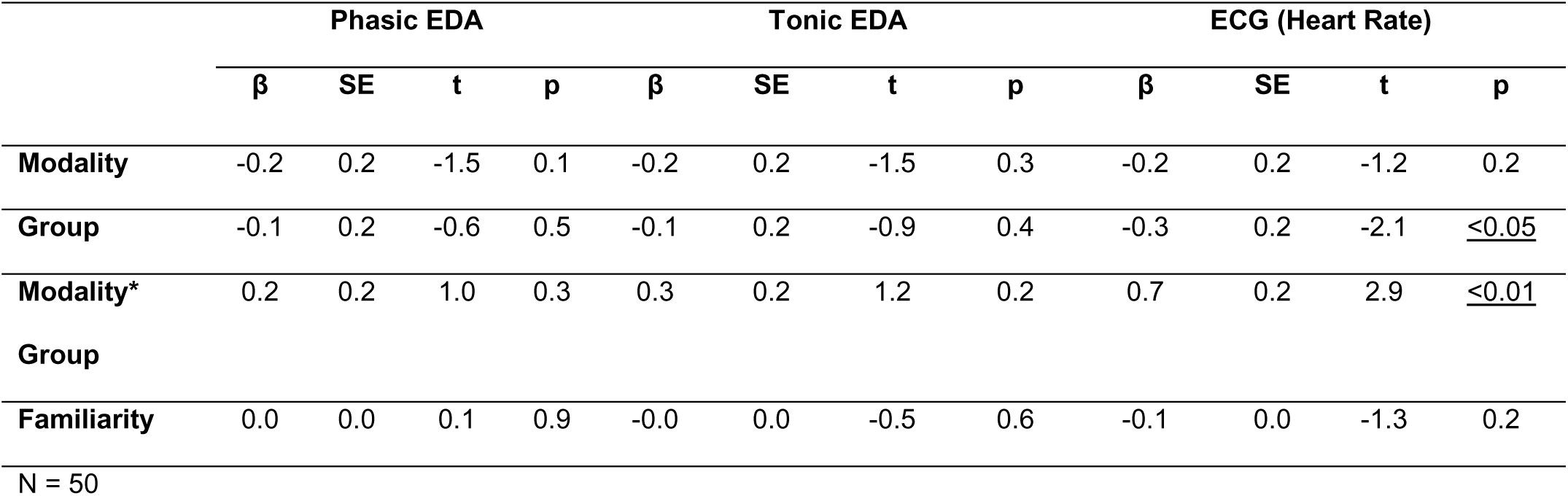
Linear mixed models predicting phasic EDA, Tonic EDA and heart rate.

##### 3.2.2.2. ECG

A significant interaction between modality and group was found (p < 0.01), with controls experiencing a higher heart rate during audio narratives compared to video narratives (β = -0.5, SE = 0.2, p < 0.05). In contrast, aphantasics had a similar heart rate while experiencing audio and video stories (β = 0.2, SE = 0.2, p = 0.6; see Table 3 and Figure 3).

**Figure 3.**
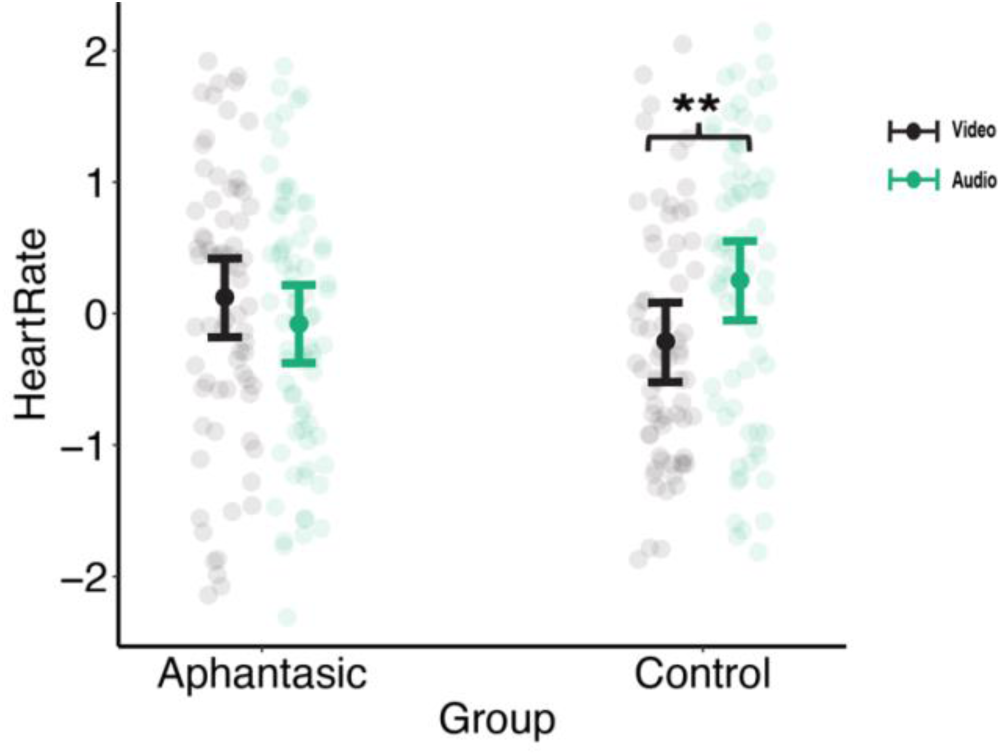
Experiment 2. Model-predicted mean and 95% confidence interval of the heart rate z-scores split between story modalities and groups. ** p<0.01

## 4. Discussion

The nature of mental imagery, along with the large individual differences in how people experience it, has been a subject of considerable debate and research, from the initial studies by Galton (1880) to arguments over whether mental imagery plays any role in cognition (Pylyshyn, 1973; Kosslyn, 1980; Shepard & Metzler, 1971; Block, 1981; Dennett, 2010). A large debate is still ongoing regarding the neural substrates supporting mental imagery (Pearson, 2019; Spagna, 2022; Dijkstra, 2024). These issues raise the question of what role visual imagery may play in everyday life. Mental Imagery has been proposed to be an “emotional amplifier’’ of thought content (Bulley et al., 2019; Chuang, 2021). Research in psychology, philosophy, and literature is rich with studies and theories delving into the interplay between mental imagery and emotional engagement in storytelling across various formats (e.g., Weibel et al., 2011; Isberner et al., 2019; Blazhenkova et al., 2023). Stories are an important part of human experience, profoundly influencing our thoughts and emotions by prompting deep reflection, impacting attitudes and beliefs, forming emotional connections, and serving as therapeutic tools for mental health (Green et al., 2003; Marsh et al., 2003; Rapp, 2016). However, studies on the role of mental imagery in narrative engagement have yielded some contradicting results and include several limitations. For example, some studies required participants to interrupt reading to report their mental imagery (Long et al., 1989; Öncel et al., 2020), while others instructed and trained participants to actively generate mental images while reading (Blazhenkova et al., 2023; Webster & Saucier, 2011; Johnson et al., 2013). Importantly, many studies only used texts expected to evoke imagery (Long et al., 1989; Johnson et al., 2013). In this study, we addressed the limitations of past research and leveraged aphantasia as a unique opportunity to investigate emotional involvement in storytelling devoid of visual imagery. Here, we use aphantasics as a ‘knockout model’ for understanding the role of mental imagery in narrative engagement in imageless cognition. In addition, we use a stimulus set that is ecologically valid, utilizing excerpts from actual movies and audiobooks. Perhaps not surprisingly, previous research suggests that canonical texts tend to be more immersive compared to stories specifically created by psychologists for experimental purposes (Green et al., 2003). Additionally, these excerpts included a wider range of emotions as they come from distinct literary genres. Lastly, considering that not all individuals with aphantasia experience a lack of auditory imagery (known as ‘Anauralia’), this study’s inclusion of audio narratives instead of readable stories provides a means to control for this potentially confounding variable, which could be advantageous for emotional engagement while reading (e.g., imagining character voices; Slámová, 2020).

Experiment 1 and 2 successfully replicated the behavioral results of Richardson and colleagues (2020): people self-report more narrative engagement (more presence, more engagement with characters, more attentional focus and more understanding) with video stories than with audio narratives (Figures 1A and 2). As in Richardson (2020), we also found an effect of familiarity, in which participants report more emotional engagement in all four subscales of the NEQ for a more familiar story. Moreover, we went one step further than Richardon and colleagues (2020) and found significant interactions of familiarity and story modality. Familiarity seems to be associated with more attentional focus and narrative presence in the case of audio, but not video, narratives (Figure 1B). As the data showed (and as many participants noted after completing the experiment), people often struggle to focus on audiobooks compared to moving-image narratives. Audiobooks are frequently used as background activity while multitasking (Nowosielski, 2018), which might explain why familiarity with a story enhances immersion, especially when distractions are minimized, as in our experiment. Audiobooks require more active interpretation and co-creation, benefiting from familiarity to fill in gaps left by the lack of visual cues. In contrast, video narratives provide rich visual stimuli that engage viewers more directly, reducing the need for familiarity to sustain attention and narrative presence.

In Experiment 2, we assessed the differences between aphantasics and controls when experiencing stories, using the same methodology as in Richardson and colleagues (2020) and as in Experiment 1. At the behavioral level, we found a significant difference between the two groups regarding the emotional aspects of narrative engagement: narrative presence and character engagement. People with aphantasia generally felt less engaged with the characters and less present in the story world compared to those without aphantasia, both when listening to audiobooks and watching videos (Figure 2). Importantly, we found no significant differences for attentional focus and understanding, which represent the more cognitive aspects of emotional engagement. This supports our original hypothesis positing that mental imagery modulates emotional but not cognitive engagement, as the results suggest that while ability to visualize did not significantly change how closely people paid attention or how well they understood the details of the narrative (whether it was in audio or video format), mental imagery did affect the way participants emotionally engaged with the narratives. That being said, it is also important to note that Dupont et al. (2023) provided evidence that individuals with aphantasia have a neurophysiological deficit in motor system engagement during action reading and struggle more than controls with deep-level comprehension. While our study found that aphantasics have the same levels of self-reported story understanding as controls, it is important to note that we did not explore the nuances of this comprehension. As Dupont et al. (2023) demonstrated, although aphantasics might grasp the phrase ‘I have a hair on my arm, I pull it out,’ they may lack the motor simulation that could deepen their understanding. This deeper understanding, which enriches the meaning of action words beyond the associative meanings processed by typical language areas, could indeed be one of the factors contributing to the heightened narrative emotional engagement observed in our sample of participants with imagery ability.

While our original hypothesis was specific to self-reported emotional engagement with audio stimuli, we observed a group-level effect that also manifested in video stimuli, indicating that the differences were not confined to audio stories alone. While it might seem intuitive to assume that imagery contributes to verbal storytelling but not to moving-image narratives, filmmakers recognize its significance, particularly in emotional scenes. They often employ techniques such as ‘Do not show the monster,’ which avoid directly showing the terrifying entity (Lyden, 2003). Instead, they build suspense and fear through shadows, character reactions, and brief glimpses, leveraging the audience’s imagination to evoke a stronger sense of fear than visual depictions could achieve. This interpretation seems especially plausible in light of research showing that aphantasics can experience fear and empathize with still images, which rules out the idea that they have a generally dampened emotional response (Wicken et al., 2021; Monzel et al., 2024). In addition, we collected data from aphantasics on their aesthetic responsiveness, primarily regarding visual arts, and found no significant differences between them and the control group. Instead, our research suggests that the emotional attenuation effect might be specific to stories. One other possibility for the observed reduced/less emotional engagement in the video modality may be specific to the exact paradigm used here and in the previous studies (Richardson et al., 2020). Before being presented with the narrative stimuli, participants read a brief introduction to the characters and plot. Presumably, this reading may have engaged mental imagery. If visual imagery helps to create a stronger sense of character and context, as hypothesized, this could also influence the amount of engagement with the characters based on this plot and character summary. In this case, aphantasics would experience less engagement with both video and audio stories because they started out less engaged when the stimulus (audio or video) appeared due to the lack of mental imagery generated during the reading of the background and plot. Recent evidence has shown that individuals with aphantasia generate content-specific representations in primary visual area (V1) when listening to evocative sounds (but not in response to instructions to voluntarily generate imagery of the sound content) despite lacking the subjective experience of imagery (Cabbai et al., 2024). Since we used audio (not readable) narratives, this might have evoked some imagery-like activity also in aphantasics, even if they could not effortfully generate it themselves. This could explain why, despite lower overall emotional engagement compared to controls, we still observed some level of emotional engagement within the aphantasic group.

Emotional engagement has been proposed as a key factor that allows stories to influence us (Quinlan & Mar, 2020; Gerrig, 1993; Gardner & Knowles, 2008). However, it’s important to recognize that engagement is not the sole pathway to enjoying a story. Our findings revealed no differences between aphantasics and controls regarding the number of novels read annually or their self-perceived proficiency as readers. Notably, one of our aphantasic participants has recently published a novel, and another is pursuing studies in literature. Additionally, Speed and colleagues (2024) showed that aphantasics did not differ in their self-reported liking or appreciation of stories than controls.

Regarding the EDA data, we were unable to replicate Richardson’s (2020) findings, which showed higher EDA during audiobook listening. We found no significant differences in either phasic or tonic EDA between audio and video stimuli and no significant effects of group or significant interactions (see Supplemental Figure 1). Our results, however, are consistent with those of Hammond et al. (2023), who also found no relationship between EDA and narrative engagement using the same equipment and the same Narrative Engagement Questionnaire assessing a group of adults sampled from the general population. To reduce the experimental session’s duration in order to minimize fatigue and boredom, we used a reduced set of stimuli compared to Richardson’s study—employing 6 trials instead of 8 (excluding the action genre stories). This omission might be one of the reasons contributing to the lack of replication, as the original effect observed by Richardson may have been primarily driven by the action stories. Additionally, a potential technical reason for not observing EDA effects could be related to the use of baseline correction methods. Wicken and colleagues (2021) found that aphantasia is associated with a flat-line physiological response to reading hypothetical frightening scenarios, which may have been due to their employment of a baseline correction method that we did not include in our methodology. This methodological difference, combined with our use of a reduced stimuli set, might have contributed to the absence of the predicted EDA effects.

In terms of heart rate, we were able to replicate Richardson’s (2020) findings only within the control group. Controls experienced a higher heart rate when listening to audiobooks compared to watching movies. However, aphantasics did not experience this difference. Richardson and colleagues (2020) argued that listening to a story requires more active imagination and co-creation than watching, making the experience more emotionally and cognitively engaging at the physiological level. This increased engagement could be driving the higher heart rate observed during audiobook sessions in their study (Richardson et al., 2020) and among controls in ours. Note that while controls behaviorally reported higher engagement for presence and character engagement than aphantasics, aphantasics (just as controls) found the videos to be more emotionally engaging than the audiobooks, suggesting that they were able to emotionally engage with the stimuli. The lack of increased heart rate in aphantasics while listening to audiobooks suggests that the increased physiological activity in controls may result from the mental effort required to generate mental imagery, rather than from emotional engagement. Cardiovascular activity is one of the most widely used physiological assessments of mental workload (Ayres et al., 2021; Tao et al., 2019; Charles & Nixon, 2019), and there is considerable evidence that mental imagery is an effortful process that can sometimes even contribute to fatigue, especially when sustained for longer periods or when involving complex or vivid imagery (e.g., Kay et al., 2022; Jacquet et al., 2021; Graham et al., 2014). This aligns with our findings, where the lack of heart rate variation in aphantasics between audio and video stories may reflect their reduced cognitive load, as they do not engage in this imagery-driven mental effort. In this context, it’s important to note the difference between mental effort (or cognitive load) and cognitive engagement. While both involve understanding and attention, mental effort refers to the degree of cognitive exertion involved. Two individuals may engage with the same content equally, yet the amount of effort each expends may differ.

All in all, our results underscore the role of mental imagery in modulating subjective and physiological responses, thereby highlighting the differential cognitive processes recruited during narrative engagement. In the future, research could delve further into the considerable variability in emotional responses among individuals with aphantasia, despite a general trend towards reduced emotional engagement within the group. As observed in our sample, some aphantasics display significantly diminished emotional reactions to stories, while others demonstrate responses that fell within the average or even above-average range. This variability highlights the need for future investigations into the factors contributing to these differences. Including hyperphantasic participants in similar experiments would be valuable, as it could facilitate clearer comparisons across the entire spectrum of mental imagery abilities. Future studies could also focus on examining the work of aphantasic writers and analyzing how their stories are perceived by both aphantasics and non-aphantasics.

## Data availability

All data can be found as supplementary materials.

## Funding

This work was supported by the Global PhD Fellowship, New York University Abu Dhabi to author N. A, and Tamkeen New York University Abu Dhabi Research Institute [grant CG012] to author D. M.

## Conflict of interest

The authors declare no conflict of interest.

## Acknowledgements

The authors thank Anna Palumbo for her assistance with ECG data analysis, Francesco Mantegna for his support with data analysis, implementation, and valuable feedback, Karleigh Groves for her help in data collection, and Maria Pombo for her assistance in implementing the online experiment.

## Notes

### Competing Interest Statement

The authors have declared no competing interest.

