## Supplemental File for "Seeing is Feeling: How Aphantasia Alters Emotional Engagement with Stories"

### Supplementary Information

#### Excerpt Synopses

Text in black was shown to participants prior to each story to contextualize the scene. Text in red is a brief summary of what the scene contained.

##### Game of Thrones

Set in the fictional Seven Kingdoms of Westeros, Game Of Thrones follows several large and wealthy families as they struggle for power in an unstable political environment. The Starks, the Baratheons and the Lannisters form an uneasy alliance - peace in Westeros balances on a knife-edge. The Baratheons and the Starks are united through an old and lasting friendship between King Robert Baratheon and Warden of the North, Ned Stark. The Baratheons and the Lannisters are united through marriage - Robert is husband to Cercei Lannister, an influential and vengeful noblewoman. Over the course of many months the uneasy truce between these three houses begins to unravel. When Ned discovers a plot against the king, learning secrets about the legitimacy of Robert and Cersei's offspring in the process, his whole family are put at risk. Now that Ned's eldest daughter Sansa is engaged to Cersei's son Jeoffrey, and his youngest daughter Arya has escaped from the Lannisters to live in disguise as a peasant, Ned must reckon with the consequences of his actions.

In the scene, Arya climbs atop a statue to observe her father’s beheading above the crowd. She sees the high lords of the realm surrounding Lord Eddard, including Joffrey and his Queen Mother. Arya’s sister, Sansa, was also on the sept amongst the lords, looking happy. Spearman holds back the crowd. Ned Stark then confesses to his treason in front of a rowdy crowd and acknowledges Joffrey as the true king. The crowd threw stones while taunting Eddard. Joffrey then orders Eddard beheaded and the executioner draws a great sword and takes Eddard’s head.

**Great Expectations**

Pip is an orphan living on the Kent marshes with his abusive sister and her husband, Joe Gargery, the village blacksmith. While exploring in the churchyard near the tombstones of his parents, Pip is accosted by an escaped convict. The convict scares Pip into stealing food for him, as well as a metal file to saw off the convict's chains. Returning with these the next morning, Pip discovers a second escaped convict, an enemy of the first one. Shortly afterward, both convicts are recaptured while fighting each other. Pip's pompous Uncle Pumblechook arranges for him to go to the house of a wealthy reclusive woman, Miss Havisham, to play with her adopted daughter, Estella. The house is strange and nightmarish.

In this scene, Pip arrives in a dressing room and meets Miss Havisham, a strange woman wearing an old bridal dress. She wants Pip to play for her entertainment. She has a sick fancy that she wants to see some play. When Pip doesn't play immediately, she calls Estella — a young woman — into the room to play cards with Pip. Estella objects to playing with the laboring boy but is forced to play regardless. Estella insults Pip until Miss Havisham invites Pip to describe his impressions of Estella and he confides that he finds her both pretty and insulting.

**SIlence of the Lambs**

Clarice Starling is pulled from her training at the FBI Academy by Jack Crawford of the FBI's Behavioral Science Unit. He assigns her to interview Hannibal Lecter, a former psychiatrist and incarcerated cannibalistic serial killer, whose insight might prove useful in the pursuit of a serial killer nicknamed Buffalo Bill, who skins his female victims' corpses. When Buffalo Bill abducts a U.S. Senator's daughter named Catherine, Jack Crawford authorizes Clarice to offer Lecter a fake deal promising a prison transfer if he provides information that helps them find Buffalo Bill and rescue Catherine.

In this scene, Hannibal offers to trade Clarice for information. He will tell her about the case if she shares information about herself. Clarice admits her worst memory was the death of her father and describes the incident. She then reminds him of their quid pro quo agreement. Hannibal asks about the girl — was she large through the hips but flat chested? She was. Clarice also admitted that the woman had an insect inserted in her throat and Hannibal asks if it was a butterfly. Clarice is surprised because this information had not been shared. Hannibal then explains what Buffalo Bill wants: he is making a lady suit from his victims.

**Alien**

Whilst exploring deep space, many, many light-years away a crew of researchers, scientists and engineers discover an earth-like planet which they decide to investigate before returning home. Having damaged their ship during the landing, a small crew breakaway from the main group and decide to explore the surrounding area for signs of habitation whilst their vessel is being repaired. During the course of their expedition, this small group of scientists come across a vast, derelict spaceship - an enormous horseshoe-shaped vessel - that looks like a promising find. As they travel deeper into the depths of this newly discovered vessel, they stumble across what remains of the ship's original crew.

In this scene, the explorers come across a room that includes eggs below a layer of mist. Soon after they realize there is movement within the eggs, the flaps atop an egg open. When the explorer takes a closer look, an alien leaps out of the egg and attacks.

#### Hound of the Baskervilles

Dr. James Mortimer asks Sherlock Holmes to investigate the death of his friend, Sir Charles Baskerville. Sir Charles was found dead on the grounds of his Devonshire estate, Baskerville Hall, and Mortimer now fears for Sir Charles' nephew and sole heir, Sir Henry Baskerville. Sir Charles' death was attributed to a heart attack, but Mortimer is suspicious, because the lord died with an expression of horror on his face.

In this scene, the client arrives to ask Sherlock and Holmes to investigate. He relates the tale of the Hound of the Baskervilles.

**Pride and Prejudice**

This tale of love and values unfolds in the class-conscious England of the late 18th century. The five Bennet sisters - including strong-willed Elizabeth and young Lydia - have been raised by their mother with one purpose in life: finding a husband. When a wealthy bachelor takes up residence in a nearby mansion, the Bennets are abuzz. Amongst the man's sophisticated circle of friends, surely there will be no shortage of suitors for the Bennet sisters. But when Elizabeth meets up with the handsome and - it would seem - snobbish Mr. Darcy, a battle of the sexes begins. Despite a testy and confrontational start to their relationship Elizabeth learns that Mr. Darcy has taken steps to save her sister, Lydia, from personal disgrace at great personal expense. Over time Elizabeth's prejudices about Mr. Darcy are challenged and she gradually comes to recognise she may have misread the wealthy nobleman.

In this scene, Elizabeth and Mr. Darcy are on a walk, chaperoned by her sister Kitty with other members of the Bennet’s walking behind. When Kitty leaves to call on a friend, Elizabeth takes her chance to thank Mr. Darcy for helping her sister. Darcy expresses surprise that the Aunt shared this information, but Elizabeth explains it was Lydia’s thoughtlessness that revealed his help. Elizabeth reiterates her thanks from her family but Darcy says, if you must thank me, let it be from you alone. He then reveals he was thinking of Elizabeth and asks her for her feelings towards him. Elizabeth re-assures Darcy that she is pleased by his feelings, making them both rather happy.

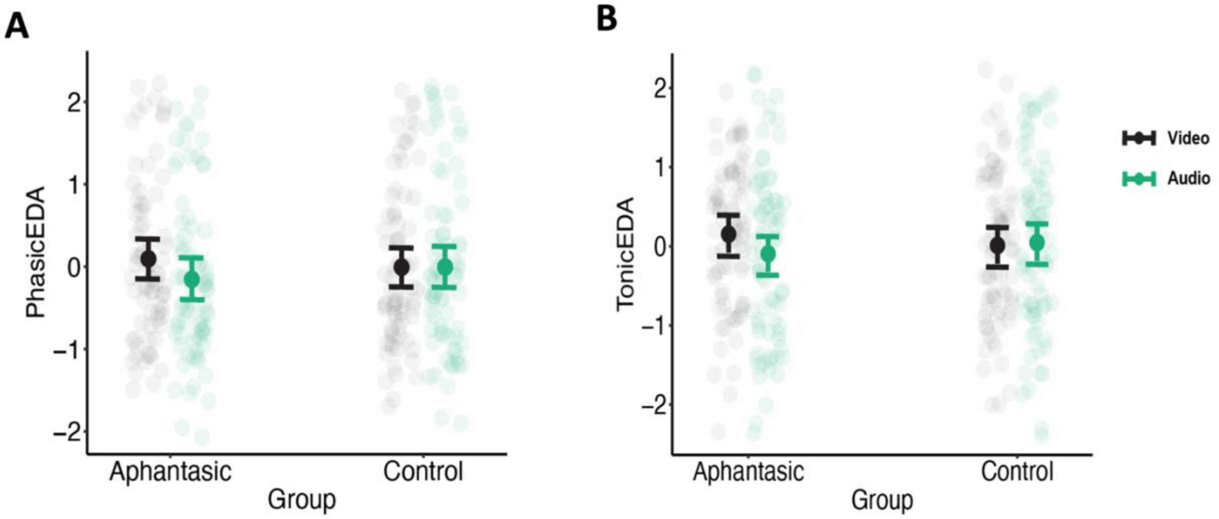


**SI Figure 1.** (A) Model-predicted mean and 95% confidence interval of the phasic EDA z-scores split between story modalities and groups. (B) Model-predicted mean and 95% confidence interval of the tonic EDA z-scores split between story modalities and groups.
